# da_Tracker: Automated workflow for high throughput single cell and single phagosome tracking in infected cells

**DOI:** 10.1101/2024.04.10.588863

**Authors:** Jacques Augenstreich, Anushka Poddar, Ashton T. Belew, Najib M El-Sayed, Volker Briken

## Abstract

Time-lapse microscopy has emerged as a crucial tool in cell biology, facilitating a deeper understanding of dynamic cellular processes. While existing tracking tools have proven effective in detecting and monitoring objects over time, the quantification of signals within these tracked objects often faces implementation constraints. In the context of infectious diseases, the quantification of signals at localized compartments within the cell and around intracellular pathogens can provide even deeper insight into the interactions between the pathogen and host cell organelles. Existing quantitative analysis at a single-phagosome level remains limited and dependent on manual tracking methods. We developed a near-fully automated workflow that performs with limited bias, high-throughput cell segmentation and quantitative tracking of both single cell and single bacterium/phagosome within multi-channel, z-stack, time-lapse confocal microscopy videos. We took advantage of the PyImageJ library to bring Fiji functionality into a Python environment and combined deep-learning-based segmentation from Cellpose with tracking algorithms from Trackmate. Our workflow provides a versatile toolkit of functions for measuring relevant signal parameters at the single-cell level (such as velocity or bacterial burden) and at the single-phagosome level (*i*.*e*. assessment of phagosome maturation over time). It’s capabilities in both single-cell and single-phagosome quantification, its flexibility and open-source nature should assist studies that aim to decipher for example the pathogenicity of bacteria and the mechanism of virulence factors that could pave the way for the development of innovative therapeutic approaches.

## Introduction

Over the past decades, the many advances in microscopy have provided fast and more precise ways to gain deeper insights into the dynamic processes of cell intracellular compartments and cells dynamics.^1^ Concurrently, the tools employed for analyzing and quantifying observed phenomena have undergone significant diversification, effectively bridging the gap between imaging advancements and the essential analysis required to quantify phenomena occurring under the microscope’s lenses.^2^

While tracking tools have proven their efficacy in detecting and monitoring objects over time, the quantification of signals within these tracked objects is often constrained to the specific image or channel used for tracking.^3^ In host-pathogen studies, single cell tracking alone cannot adequately analyze the dynamics of intracellular bacteria within host cells. Studies often employ up to four markers including the pathogen, to comprehensively analyze these interactions, to track the host cell response and the localization and dynamics of the pathogen. Thus, there is a compelling need to integrate cell tracking workflows with flexible quantification methodologies. In the context of mycobacterial studies, time-lapse microscopy was used to dissect the dynamics of the mycobacteria containing phagosome (MCV), like its maturation, damages, interactions with the host autophagy, or the bacteria growth.^4–12^ In this context, several methods were proposed to track and quantify recruitment of factors at the MCV from time-lapse confocal imaging datasets.^13–15^

While some of these methods provide robust workflows to study phagosome maturation or intracellular growth, the cell tracking relies on manual cropping of the cell and the bacteria tracking is mostly performed at least partially manually in the case of confocal microscopy, or relies on proprietary software.

The main limitation in existing tracking tools is that they don’t necessarily provide manipulatable data structures which allow expanded downstream analyses. In the ImageJ environment the most first candidate is the Region Of Interest (ROI); which is flexible and provides a powerful capability for precisely isolating intracellular signals. Indeed, the delineated ROIs generated by segmentation software not only encompass the cell boundaries but also contains the entirety of the signal from adjacent slices, especially in the context of a cell monolayer. This allows the quantification of signals of interest emanating from different intracellular compartments or objects, facilitating the subsequent tracking of these signals throughout the duration of a time-lapse experiment. This becomes particularly interesting in scenarios involving infected cells hosting bacteria or similar intracellular objects of interest. By directly tracking and quantifying markers association with either the bacteria or the phagosome membrane, it becomes feasible to analyze and quantify in high throughput the intricate interplay between host cell defense mechanisms and the pathogenic bacteria’s evasion mechanisms.

In the absence of an available microscopy system allowing the physical separation of cells, the cells segmentation can be accomplished using various available software. The segmentation software Cellpose^16,17^ offers a unique feature, enabling the easy training of custom models to better fit specific samples as demonstrated in some of our previous work on cells or bacteria.^18,19^ Morphological disparities between mouse and human cells, or even amoebae could potentially hinder accurate cell recognition by a trained model; however, Cellpose addresses this challenge by allowing flexible retraining of models to meet the demands of individual experiments. In terms of quantification and versatility, ImageJ / Fiji serves as a widely used and robust platform, particularly through the use of ROIs, which facilitate the transfer of segmentation results onto any image for subsequent signal quantification and data collection. Moreover, Fiji comes equipped with numerous built-in plugins, including for tracking purposes, and thus offers diverse analytical possibilities. In addition, PyimageJ library allows the manipulation of ImageJ/Fiji inside the Python ecosystem,^20^ enabling interoperability between ImageJ and different Python-based software like Cellpose in the same environment. This interoperability has demonstrated its potential to significantly enhance automation and batch processing capabilities.^18^ Although segmentation software like Cellpose can eventually return ROIs, it does not perform tracking calculation. A recent extension of the Fiji tracking plugin Trackmate^21^ integrated Cellpose as a cell detector, thus bridging the gap between segmentation and tracking, allowing the extraction of ROIs as well.

In this article, we present an advanced image analysis workflow that combines deep learning-based cell segmentation using Cellpose with the powerful quantification and flexibility of ImageJ/Fiji and Trackmate, integrated within the Python platform PyimageJ. This novel approach allows to perform single cell and single bacterium/phagosome tracking and fluorescence quantification in a high-throughput fashion. It also provides a free and open-source solution for studying host-pathogen interactions and other related biological phenomena.

## Results

### Workflow for single cell and single phagosome tracking

This workflow is designed to provide a comprehensive and near-fully automated solution for the quantitative tracking of single cells and bacteria within multi-channel, z-stack time-lapse images, followed by the single bacterium / phagosome tracking often applied in the context of infections and similar studies. The entire workflow is visually represented in a diagram in Figure 1. The source code can be found on GitHub (URL: [notebook](https://github.com/jaugenst/da_tracker/blob/main/notebook/da_tracker_notebook.ipynb)) as a Jupyter notebook.

**Figure 1:**
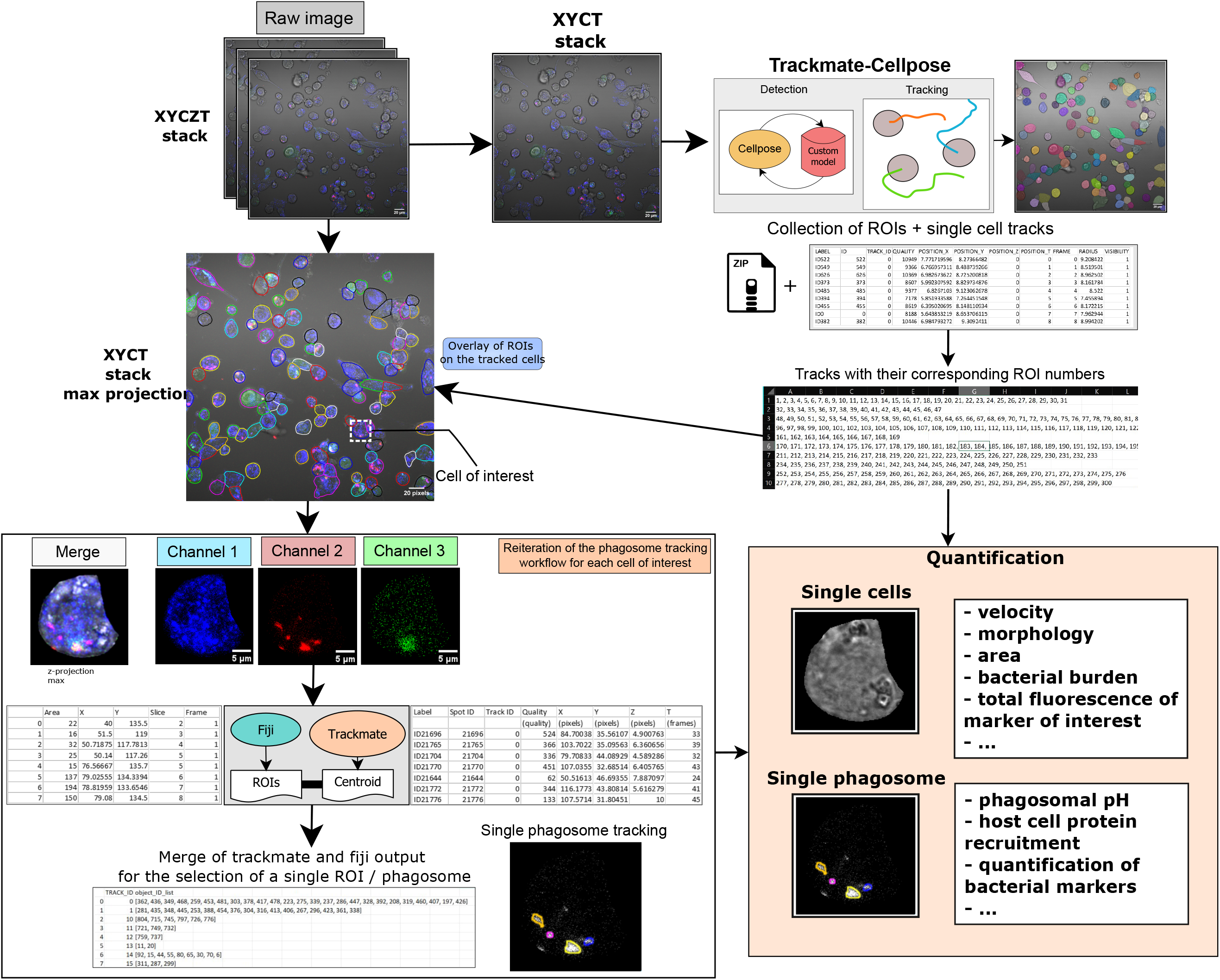
Diagram of the workflow for quantitative single cells and single bacteria tracking. Diagram of the workflow with representative microscopy images from each step. The “table output” and “plot” are representative of the quantification process.

#### Step 1: Data Extraction and Preparation

The workflow begins by extracting three sets of images from the raw time-lapse file. First it extracts a single z, multi-channel time-lapse that would allow Trackmate-Cellpose to segment and track the cells from the T-PMT channel. In the current implementation, the process of extracting ROIs from a single z, single channel time-lapse does not bind the resulting ROIs to the time frame. Instead, they are bound to the first frame, preventing any extraction of signals of the cells over time. In order to avoid this, the image must be a multi-channel and/or multi-z. The next image is a z-projection ‘max’ of the raw file that would be used for the curation of the cells’ trajectory. All the images are saved into the main directory. The different channels are also isolated as single channel, z-stacks time-lapses. This is required for the downstream single cell signal isolation.

#### Step 2: Cell segmentation and tracking using Trackmate-Cellpose

Deep learning-based segmentation has the potential to detect and segment cells where classical thresholding approaches are insufficient. Cellpose offers a flexible environment to train and segment images at will and adapt the model to the samples presenting different morphologies. The process of training of a custom Cellpose model is already described in the associated published work and documentation.^16^ Trackmate is a tracking plugin integrated in Fiji that recently incorporated Cellpose as a detector and Trackmate’s use is also well documented.^21^ The Trackmate-Cellpose plugin allows the user to call a specified custom model or pre-trained model. The plugin provides a python API, allowing full automation if the detection parameters are already known and adjusted. After the segmentation using Cellpose in Trackmate, the tracking is done according to the documentation. It is to be noted that before using the script in PyimageJ, it is first recommended to use a time-lapse from a representative field of view to define the different parameters of tracking (e.g.: algorithm, quality and threshold values) in the Trackmate interface itself. Indeed, the interface is optimized to change some parameters and see the results of the change very efficiently. The detection and the tracking parameters can be modified in a dedicated code block in the notebook. The tracking output is saved in the main directory.

#### Step 3: Trackmate output treatment and ROI extraction

Once the tracking is performed, a CSV file named ‘exportTracks’ is exported containing all the parameters of interest for each detected cell at one time point or ‘spots’ in the Trackmate table output. The trajectories correspond to the ‘Tracks’ and their associated number are in their designated column. From this table, the spots in each track are first sorted in chronological order and saved under the name ‘exportTracks_edited’. Then, the index of each spot for each track is stored in a new csv table named ‘da_tracks’. In the latter, each lane will contain a list of indexes corresponding to each track or cell. Then, the ROIs are open in the ROI manager sequentially for each spot of each track in chronological order using the spot ‘LABEL’. Ultimately the ROI manager is populated with all the ROIs from Cellpose, chronologically sorted and match their respective tracks and spots.

#### Step 4: Cell trajectory curation and visualization

For each track, the associated ROIs are selected and overlayed on the z-projection ‘max’ image saved in step 1, with a randomly picked color for each track. This allows the user to visually assess the quality of the trajectory and if it was not doable in the Trackmate tracking settings, to correct and manually refine the trajectories. Indeed, by using the ‘show labels’ option in the ROI manager, the cells also display a number, that will match the ROI number lists in the file ‘da_tracks.csv’. This file may be modified to manually correct detected trajectories or choose a specific subset of interest (i.e. infected cells) or to exclude cells detected for an insufficient time period. For this selection, the user would first have to retain the first number that appears on the cell at the start of its trajectory, then find the corresponding line number in the da_tracks file. This number correspond to a cell number that can be added to the list ‘cells_desired’ in the corresponding code block. In case the trajectory is fragmented into 2 or more tracks, the user can also add a list of rows into the ‘cells_desired’ list.

#### Step 5: Single cell time lapse isolation

To efficiently detect and track a single phagosome in a single cell, the signal from extracellular bacteria needs to be excluded. As the cells are also mobile, to decrease the frame-to-frame displacement of the phagosomes, the signal from the cell is also isolated and reassembled as a single time-lapse. To do so, for each cell number in the ‘cells_desired’ list, the ROIs are selected one by one, the cell area is duplicated and the signal outside the area is deleted. This image is then concatenated in a new time lapse. The result for each cell is a time-lapse of z-stacks centered around the single cell. The process is reiterated for each channel of interest for further analysis in addition to the bacterial channel. The latter is saved in a dedicated directory named ‘bact’ while the other channels are saved in a directory named ‘other’. Ultimately, this clean intracellular signal can now be used to perform the detection and tracking of the phagosome.

#### Step 6: Single bacterium / phagosome detection and tracking

From the single cell bacterium channel saved in step 5, the bacteria are first detected in Fiji using the more classic approach of thresholding followed by particle detection to obtain a set of ROIs for each slice of each bacterium combined with the measurement of their XY coordinates. The bacterial channel is also analyzed in Trackmate, using the threshold detector and the trajectories calculated with the LAP tracker. The objective is to collect the bacterium centroid coordinates contained in the Trackmate ‘spots’ table output. It is to be noted that the parameters given in the code were optimal for our samples but the detector type and tracking parameters can be fine-tuned to fit the user sample. Finally, the bacterium centroid coordinates from Trackmate were compared to the XY coordinates of the ROIs obtained from the particle detection, using the nearest neighbor calculation from the geopandas library in Python. The closest ROI from the bacterium centroid is retained and the calculation is reiterated for each frame. The ROI number list matching each bacterium trajectory is stored in a new file similar in structure of the previous ‘da_tracks’ file, where each row contains the list of ROIs of an individual tracked bacterium. The file is in a subdirectory named ‘grouped’ in the ‘bact’ directory created in step 5 and is named after the cell number in the ‘cell_desired’ list. An additional quality control step of tracking quality is performed by overlaying every trackmate-derived ROI on the appropriate slice of the bacterial channel. The image is then flattened, compressed as a z-projection ‘max’, and saved with ‘overlay’ as an extension to the name. Each bacterial track would be with overlayed with a randomly picked color thus allowing to visualize and evaluate the quality of the tracking results, in the same fashion as the step 4.

#### Step 7a: Measurement of parameters of interest at the single cell level

The extant workflow provides examples illustrating the measurement of parameters of interest on single cells across channels. Thus, it is possible to describe various aspects of the data including cell velocity, cell size change bacterial burden, or relative abundance or markers by the measure of the total cell fluorescence on the z-projection ‘sum’ generated in step 1. These outcomes can be obtained by running the provided code located in the ‘Single cell quantification’ section of the notebook. The tables containing the final measurements are saved in a directory called ‘misc’.

#### Step 7b: Measurement of parameters of interest at the single bacteria level

The fluorescence measurements of interest can be collected on the bacteria in the other channels and the workflow provides a code block where the user can give the names of the different channels of interest. Given the desired channels, the extant workflow opens them sequentially, applies the detected ROIs, and measures the parameter of interest, e.g. the mean fluorescence intensity. This is repeated for each phagosome’s trajectory and each channel. The measurement output is given as a CSV file containing a table combining the fluorescence values from the channels of interest chronologically ordered and sorted by their respective track. This outcome can be obtained by running the provided code located in the ‘Single bacteria feature measurement’ section of the notebook. The tables containing the final measurements are saved in a directory called ‘signal measurement’.

### Example of results

This workflow was originally designed in the context of THP-1 cells infected with *Mycobacterium tuberculosis* (Mtb). However, in our current microscopy configuration, the mobility of differentiated THP-1 cells was too great (Movie S1) and attempts to further increase the frame rate would have been made at the expense of z-stack range or number of cells imaged in one experiment. We then tested the workflow on primary human Monocytes-Derived Macrophages (hMDMs) infected with Mtb. The results revealed a slower and more regular motion of hMDMs (Movie S2), more suitable for optimal tracking of cells and Mtb-containing vacuoles (MCVs) (Movie S3). As described above, the cells of interest in this case cells infected with1-3 MCVs were selected. After the single cells signals were isolated and collected, the bacteria tracking workflow was applied. The results in figure 2 show an example of the metrics and data that can be collected from the entire workflow. We separated the data collection into 2 categories in the notebook, the single cell level and the single bacterium / phagosome level. For the single cell level, the collection of the single cell positions in the image allows the calculation of the cell velocity, while the outline of the cells allows the collection of the area value over time (Figure 2A). As we also released a method to measure the relative bacterial burden,^18^ the code was adapted to calculate the bacterial volume directly from the bacterial ROI collection in step 6 and was extended over time. The bacterial volume over time was calculated for each single cells (gray curves) and the overall average can also be visualized (Figure 2A). This curve can then be fitted with, for example, a linear regression to estimate the growth rate in this hMDMs over ∼30h. Finally, another example of a readout is the measurement of total fluorescence of lysoview which should reflect the relative quantity of lysosomes and late endosomes (Figure 2A).

**Figure 2:**
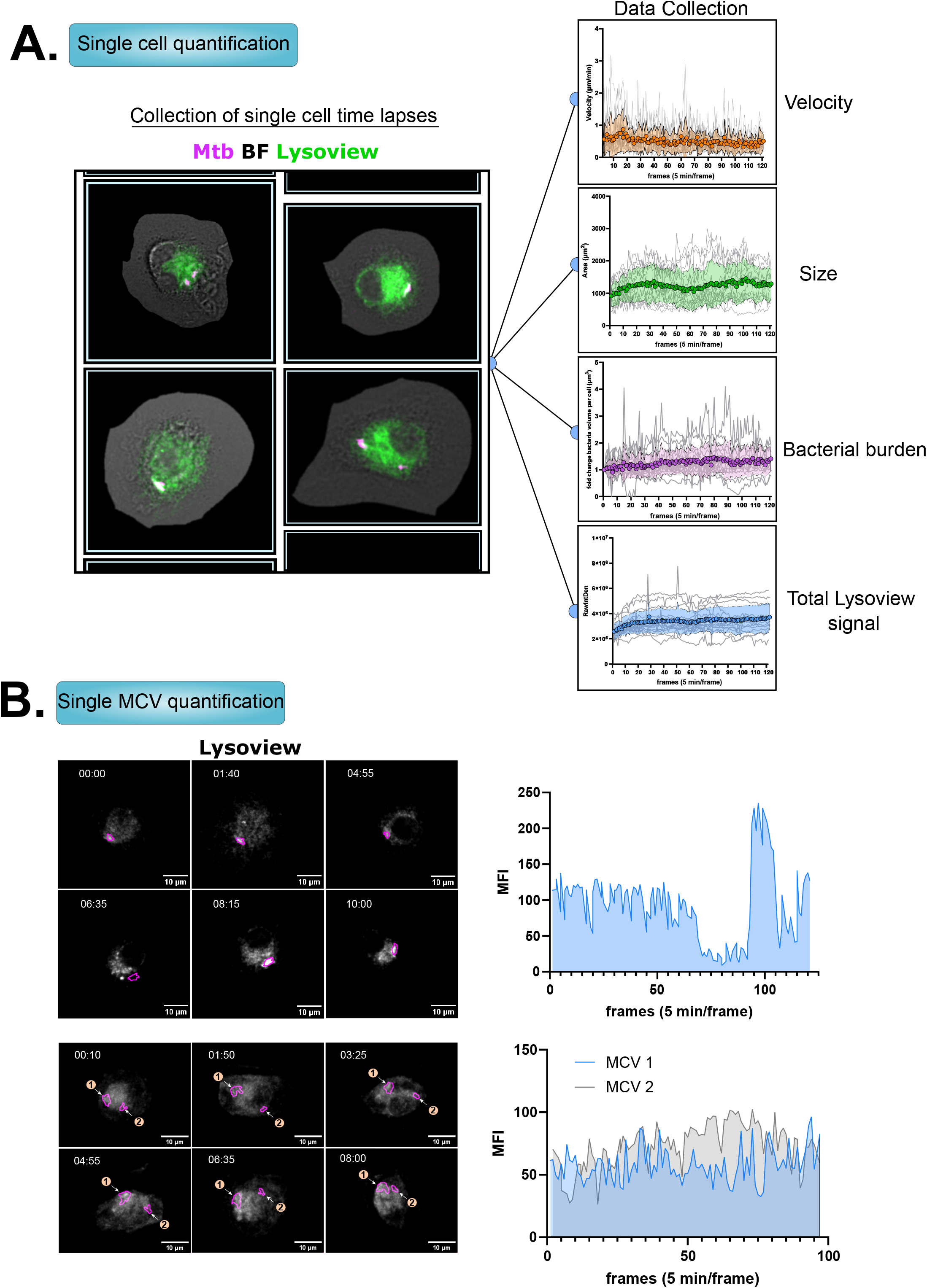
Quantification of single cells features and single bacteria fluorescence in infected hMDMs. (A) Left: representative images of individual single cells’ time-lapses selected from Movie S3. Right: Quantification of cells’ velocity, area, bacterial burden and total lysoview-633 fluorescence over time. The individual curves in gray correspond to the single cell quantification. The colored curve with transparent band represents the mean and standard deviation of the single curves. (B) Left: Representative time-lapse images from the lysoview-633 channel of cells bearing one (top, Movie S4) or two MCVs (bottom, Movie S5). Right: quantification of the mean fluorescence intensity (MFI) of lysoview-633 in the magenta area in the time-lapse.

At the MCV level, from the single cell signal the bacteria were tracked following the workflow. The example here is given for cells with one or two MCVs with a tracking result showing a continuous monitoring of the MCV for more than 16h (Figure 2B, Movie S4-5). The lysoview MFI was quantified over time and demonstrates the potential of the workflow to quantify signals in or around a bacterium from long term time-lapse acquisition (Figure 2B).

The provided code gives a highly automated way of collecting these numbers in minutes after the tracking was performed. The use of PyimageJ in python allows for adaptability and the provided code can be modified for other classical analysis readouts. It could also be combined with existing methods to analyze phagosome maturation^13,14^ and could be automatized for a time lapse. The application for this workflow could range from the studies of association of proteins to the bacterial phagosome, to the monitoring of bacterial burden during drug treatment tests. While not tested, the workflow should also be theoretically applicable to any bacteria of interest, and in different host cell systems.

One limitation of this workflow is in analysing highly mobile cells because that can lead to cells leaving the field of view. Another limitation came from the microscope itself (Zeiss LSM-800) since we observed sub-optimal tracking result if the frame rate is too low on mobile cells. For example, in our experiments a frame rate of 10 min resulted in fragmented tracking (data not shown), and thus we increased the frame rate to 5 min. However, increasing the frame rate on a classical confocal microscope configuration is to be done at the expense of the number of fields of view, the range or the interval of the z-stack. For the latter, a slice sampling too spaced or a too narrow range could induce temporary loss of bacterial signal and fragmented tracking. The current perspectives to address these issues are to apply this workflow on datasets obtained from high-speed confocal microscopy techniques, such as spinning disk microscope, or lattice light-sheet microscope. These techniques are also known to be gentler in terms of illumination of the sample, thus may prove optimal for the proposed high throughput time-lapse analysis of infected cells in high resolution as well.

In conclusion, this workflow presents a practical automated avenue for conducting quantitative single cell tracking followed by single phagosome tracking and fluorescence analysis, through the PyimageJ processing capabilities in combination of cell segmentation, trajectory analysis, and multi-metric measurements. This is done using free and open-access tools thus that can be used by anyone having access to microscopy equipment. This integrated workflow should be a potent toolkit for single cell and single phagosome tracking. Its adaptability, and time efficiency should be of great help in the context of cellular research and host-pathogen interactions.

## Material and methods

### Software and installation

The code and process are described in detail in the associate notebook available on GitHub (https://github.com/jaugenst/da_tracker).

It is highly recommended to install any software and libraries required for this workflow in a virtual environment, using Anaconda, Pyvenv or environment modules. A detailed documentation of PyimageJ is available online (https://py.imagej.net/en/latest/). The workflow development was done in a jupyter notebook in the software jupyter-lab (https://jupyter.org/). Cellpose software was installed following the documentation and installation instructions available online (https://github.com/mouseland/cellpose). Trackmate-cellpose was installed in Fiji following the available documentation (https://imagej.net/plugins/trackmate/detectors/trackmate-cellpose). ChatGPT 3.5 (OpenAI) was occasionally used for code debugging, editing and text editing.

### Reagents

Phorbol-Myristate-Acetate (PMA, provider), RPMI medium (Gibco), Fetal bovine serum (thermofisher), human off-the-clot serum (Geminibio), human recombinant macrophage-colony stimulating factor (M-CSF, R&D Systems), Lysoview-633 (biotium), OADC supplement (R&D Systems), Zeocin (invivogen).

### Cell culture, bacterial culture

GFP-tagged LC3 expressing THP1 monocytes were cultured as previously described (ref). The cells were cultured in RPMI 1640 (ATCC modification) supplemented with 10% heat-inactivated Fetal bovine serum. The cells were differentiated with 50 ng/ml of PMA for 48 hours and allowed to rest in complete medium for 48h before the infection.

Peripheral blood mononuclear cells (PBMCs) from healthy blood donors were isolated from leukopaks by using Ficoll density gradient centrifugation. For monocyte isolation, 1x10^6^ PDMCs were seeded in an 8-well ibidi uncoated-bottom chamber for 2 hours in unsupplemented RPMI. The attached monocytes were washed and incubated for 7 days in RPMI 1640 supplemented with human off-the-lot serum 5% and M-CSF at 10ng/mL (complete medium).

*Mycobacterium tuberculosis* H37Rv (ATCC) expressing DsRed were grown in liquid Middlebrook 7H9 medium supplemented with 10% oleic acid-albumin-dextrose catalase (OADC) growth supplement, 0.2% glycerol, 0.05% tween 80, and Zeocin 100µg/mL. For infections, the bacteria were washed in 0.05% PBS-tween 80 before being added to the infection medium (see procedure below).

### Cell infection

The bacteria were pelleted and washed in PBS-tween80 0.05%. The OD600 was measured, and the bacteria were added on the hMDMs at an MOI of 2 for 1 hour. The cells were washed three times with unsupplemented RPMI and incubated with fresh complete medium supplemented with lysoview-633 diluted at 1:2000.

### Time-lapse confocal imaging

The cells were imaged using a Zeiss laser scanning confocal microscope LSM-800, equipped with two gallium arsenide phosphide photomultiplier tube (GaAsP-PMT) detectors and a transmitted light photomultiplier tube detector (T-PMT), using the 40x/NA1.2 water objective. The observation chamber was maintained at 37°C and supplied with 5% CO2 through a humidifier system. A 10 μm z-stack (1 μm steps) was acquired every 5 minutes on 2 fields of view over 10 hours.

### Data analysis, and data visualization

The graphics were generated using GraphPad prism 10. All the figures and drawings were made and assembled in Inkscape.

## Author contributions

Conceptualization: J.A., V.B.; Methodology: J.A., A.T.B., A.P.; Software: J.A., A.T.B.; Validation: J.A., A.T.B., A.P.; Formal analysis: J.A.; Investigation: J.A.; Resources: J.A., A.T.B., A.P.; Data curation: J.A.; Writing - original draft: J.A.; Writing - review & editing: J.A., A.T.B., V.B., N.E.-S.; Visualization: J.A.; Supervision: J.A., V.B.; Project administration: J.A., V.B.; Funding acquisition: V.B.

## Data availability

The code used for the workflow and the cellpose models for segmentation are provided on GitHub (https://github.com/jaugenst/da_tracker/notebook/). The supplemental movies can be found on Mendeley data here^22^ or can be obtained using the following download link:

Movie S1, Movie S2, Movie S3, Movie S4, Movie S5

## Acknowledgments

We sincerely thank Dr Serge Mazeres his constructive feedback for the workflow development.

## Figure Legends

**Movie S1:** THP-1 cells expressing LC3-GFP were infected with H37Rv-DsRed at an MOI 2 for 1h and imaged for 10h. A 10μm z-stack was acquired every 5 minutes. Time stamp format is hh:mm, and only one slice of the transmitted light channel is shown.

**Movie S2:** hMDMs were infected with H37Rv-DsRed at an MOI 2 for 1h and imaged for 10h. A 10μm z-stack was acquired every 5 minutes. Time stamp format is hh:mm, and only one slice and the transmitted light channel are shown.

**Movie S3:** hMDMs were infected with H37Rv-DsRed (magenta) at an MOI 2 for 1h, stained with Lysoview-633 (green) and imaged for 10h. This movie is the same field of view than the movie S2. The transmitted light channel is displayed in Gray. A 10μm z-stack was acquired every 5 minutes. Time stamp format is hh:mm. The movie is a z-projection ‘max’ of the original time-lapse. Each cell detected by the tracking workflow is outlined with a randomly picked color.

**Movie S4:** Example tracking a single MCV (magenta outline) in the lysoview channel (gray) over time. Time stamp format is hh:mm. The movie is a z-projection ‘max’ of the original time-lapse.

**Movie S5:** Example tracking two MCVs (magenta outlines) in the lysoview channel (gray) over time. Time stamp format is hh:mm. The movie is a z-projection ‘max’ of the original time-lapse.

## References

1. Ouyang, W. & Zimmer, C. The imaging tsunami: Computational opportunities and challenges. Curr. Opin. Syst. Biol. 4, 105–113 (2017).

2. Haase, R. et al. A Hitchhiker’s guide through the bio-image analysis software universe. FEBS Lett. 596, 2472–2485 (2022).

3. Caicedo, J. C. et al. Data-analysis strategies for image-based cell profiling. Nat. Methods 14, 849–863 (2017).

4. Mahamed, D. et al. Intracellular growth of Mycobacterium tuberculosis after macrophage cell death leads to serial killing of host cells. eLife 6, e22028 (2017).

5. Schnettger, L. et al. A Rab20-Dependent Membrane Trafficking Pathway Controls M. tuberculosis Replication by Regulating Phagosome Spaciousness and Integrity. Cell Host Microbe 21, 619-628.e5 (2017).

6. Lerner, T. R. et al. Mycobacterium tuberculosis replicates within necrotic human macrophages. J. Cell Biol. 216, 583–594 (2017).

7. Cardenal-Muñoz, E. et al. Mycobacterium marinum antagonistically induces an autophagic response while repressing the autophagic flux in a TORC1- and ESX-1-dependent manner. PLOS Pathog. 13, e1006344 (2017).

8. López-Jiménez, A. T. et al. The ESCRT and autophagy machineries cooperate to repair ESX-1-dependent damage at the Mycobacterium-containing vacuole but have opposite impact on containing the infection. PLOS Pathog. 14, e1007501 (2018).

9. Bernard, E. M. et al. M. tuberculosis infection of human iPSC-derived macrophages reveals complex membrane dynamics during xenophagy evasion. J. Cell Sci. 134, jcs252973 (2020).

10. Beckwith, K. S. et al. Plasma membrane damage causes NLRP3 activation and pyroptosis during Mycobacterium tuberculosis infection. Nat. Commun. 11, 2270 (2020).

11. Rutschmann, O., Toniolo, C. & McKinney, J. D. Preexisting Heterogeneity of Inducible Nitric Oxide Synthase Expression Drives Differential Growth of Mycobacterium tuberculosis in Macrophages. mBio 0, e02251–22 (2022).

12. Raykov, L., Mottet, M., Nitschke, J. & Soldati, T. A TRAF-like E3 ubiquitin ligase TrafE coordinates ESCRT and autophagy in endolysosomal damage response and cell-autonomous immunity to Mycobacterium marinum. eLife 12, e85727 (2023).

13. Barisch, C., López-Jiménez, A. T. & Soldati, T. Live Imaging of Mycobacterium marinum Infection in Dictyostelium discoideum. in Mycobacteria Protocols (eds. Parish, T. & Roberts, D. M.) 369–385 (Springer, New York, NY, 2015). doi:10.1007/978-1-4939-2450-9_23.

14. Schnettger, L. & Gutierrez, M. G. Quantitative Spatiotemporal Analysis of Phagosome Maturation in Live Cells. in Phagocytosis and Phagosomes: Methods and Protocols (ed. Botelho, R.) 169–184 (Springer, New York, NY, 2017). doi:10.1007/978-1-4939-6581-6_11.

15. Arévalo, P. R., Aylan, B. & Gutierrez, M. G. Quantitative Spatio-temporal Analysis of Phagosome Maturation in Live Cells. in Phagocytosis and Phagosomes: Methods and Protocols (ed. Botelho, R. J.) 187–207 (Springer US, New York, NY, 2023). doi:10.1007/978-1-0716-3338-0_13.

16. Stringer, C., Wang, T., Michaelos, M. & Pachitariu, M. Cellpose: a generalist algorithm for cellular segmentation. Nat. Methods 18, 100–106 (2021).

17. Pachitariu, M. & Stringer, C. Cellpose 2.0: how to train your own model. Nat. Methods 19, 1634–1641 (2022).

18. Augenstreich, J. et al. BBQ methods: streamlined workflows for bacterial burden quantification in infected cells by confocal microscopy. Biol. Open 13, bio060189 (2024).

19. Lyu, Z. et al. Genome-wide screening reveals metabolic regulation of stop-codon readthrough by cyclic AMP. Nucleic Acids Res. gkad725 (2023) doi:10.1093/nar/gkad725.

20. Rueden, C. T. et al. PyImageJ: A library for integrating ImageJ and Python. Nat. Methods 1–2 (2022) doi:10.1038/s41592-022-01655-4.

21. Ershov, D. et al. TrackMate 7: integrating state-of-the-art segmentation algorithms into tracking pipelines. Nat. Methods 19, 829–832 (2022).

22. Augenstreich, J. & Briken, V. Supplemental movies for da_tracker workflow. 1, (2024).

